# TOG domain MT polymerases accelerate MT plus end growth via electrostatically-steered diffusion-to-capture and electrostatic templating of GTP-tubulin

**DOI:** 10.1101/282897

**Authors:** Neil A. Venables, Till Bretschneider, Robert A. Cross

**Affiliations:** Centre for Mechanochemical Cell Biology, Warwick Medical School, Gibbet Hill, Coventry CV4 7AL; Department of Computer Science, University of Warwick, Coventry, CV4 7AL

**Keywords:** TOG domains, Electrostatics, Microtubules, Brownian Dynamics, Molecular Dynamics, Tubulin

## Abstract

TOG domain microtubule polymerases track microtubule plus ends, bind GTP-tubulin and catalyse microtubule growth, by mechanisms that are not yet understood. In this work, we use computational analysis and simulation to probe the detailed mechanism of tubulin capture and exchange by TOG domains. TOG domains display a ridge of 5 surface loops that form the core of the TOG-tubulin interface. Using computational mutagenesis, we confirm that this row of loops, which is positively charged, plays a dominant role in setting the overall electrostatic field on the TOG domain. Brownian dynamics simulations establish that diffusion-to-capture of TOGs by tubulin is very strongly electrostatically steered. Under a range of conditions and in all trajectories examined, TOGs are initially captured and oriented by tubulin at high radius so that their basic loops faced inwards towards the tubulin. Thereafter, the loops continue to face inwards towards the tubulin and to scan its surface until stereospecific docking the crystallographic binding site occurs. We find that the acidic C-terminal tails of tubulin are not required for electrostatic steering, but instead serve to widen the acceptance angles for electrostatically steered diffusion-to-capture. All-atom normal mode analysis indicates that TOGs are remarkably stiff, enabling them to drive free GTP-tubulin into a partially-curved state by conformational selection. Electrostatic free energy calculations show that the complex that each TOG makes with its cognate tubulin is stable. Our work argues that TOGs accelerate microtubule plus end growth by two complementary electrostatic mechanisms, first by electrostatically steered diffusion-to-capture, and second by electrostatic stabilisation of a partially bent conformation of GTP-tubulin that exchanges rapidly into the tip-lattice. To explain this rapid exchange, we propose a model in which simultaneous binding of the GTP-tubulin to the TOG and the microtubule tip-lattice can occur and is required to de-stabilise the crystallographic complex and release and recycle the TOG.

**Author Summary:** TOG domain microtubule polymerases are protein machines that accelerate the growth of microtubule plus ends by capturing tubulin building blocks from solution and feeding them into the growing microtubule tip. Exactly how TOGs manage to do this remains unclear. Several lines of evidence suggest that electrostatic interactions play a key role, but the detailed role of electrostatics in the polymerase mechanism of TOGs is so far little explored. Here using linked computational approaches we analyse the electrostatic fields of TOGs from the TOG polymerase superfamily and simulate their tubulin binding trajectories. We find that each TOG domain has a shaped electrostatic field that is precisely matched to its tubulin binding partner, such that each TOG is electrostatically orientated at high radius and thereafter electrostatically guided to its capture site. Our work shows that electrostatic steering dramatically accelerates the diffusion-to-capture of tubulin by TOGs. The resulting TOG-tubulin complexes are electrostatically stabilized and we suggest that release of the TOG from this complex requires that tubulin first be incorporated into the growing microtubule, thereby being driven into a TOG-incompatible conformation.

## Introduction

TOG domain MT polymerases track the growing plus-ends of dynamic microtubules (MTs) and accelerate their growth by around 10-fold [1]. There are now several crystal structures of TOG-tubulin complexes [2, 3], that reveal their detailed structure. However the dynamic mechanisms of GTP-tubulin exchange by TOG domains, and their relationship to the modulation of MT growth, remain poorly understood.

TOG domains share a common architecture in which six HEAT-repeats (HRs) [4, 5] pack side-by-side to form a paddle-like structure [6]. Each HEAT repeat is composed of two antiparallel alpha helices connected by a loop. In the TOG domain, several intra-HEAT loops align to form a positively charged ridge (see **Fig. 1 A**) that fits into a negatively-charged channel on the tubulin dimer (see **Fig. 1 B**). TOGs preferentially bind to unpolymerised tubulin in a nucleotide independent manner (see **Fig. 1 C**) [2]. Mutation of conserved residues in the intra-repeat turns disrupts tubulin binding [6].

**Fig 1.**
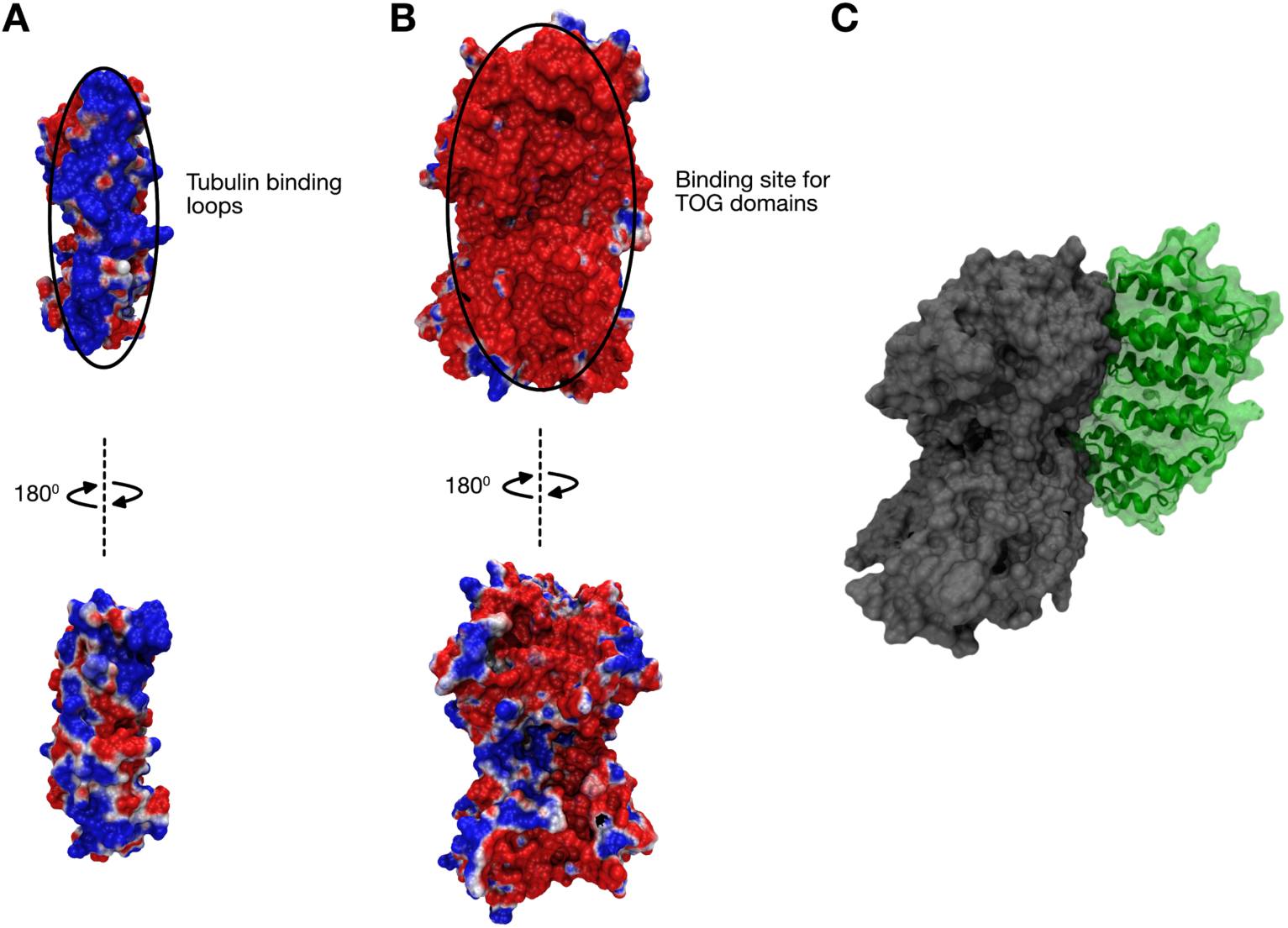
Complementary electrostatic surfaces of the TOG domain and tubulin dimer. Electrostatic surface projection for both the TOG2 domain (A) and tubulin dimer without C-termini (B) from *S. cerevisiae* (PDB: 4U3J). Structures are coloured along a gradient from −1 k_b_T/e (red) to +1 k_b_T/e (blue), and show complementary charges in their respective binding regions. Below, each structure is rotated 180°, showing a heterogeneous charge landscape away from the putative binding site. (C) Surface representation showing how the binding regions interact to form a stereospecific complex with the tubulin in grey and the TOG domain in green.

MT assembly is a complex and diffusion-limited process [7]. The apparent on-rate constant for GTP-tubulin at growing MT plus ends was measured at 52 µM^-1^ s^-1^ for porcine brain tubulin *in vitro* [8]. This rate constant is consistent with the theoretical limit calculated for a diffusion-controlled reaction of around ∼4 µM^-1^ s^-1^ per MT protofilament [9]. TOG-family polymerases can increase the MT net growth rate at plus ends by up to ∼10-fold. Several lines of evidence suggest that electrostatic interactions play a key role in TOG-tubulin interactions [10, 11], but the detailed role of electrostatic interactions in determining the dynamics of complex formation and / or the catalysis of tubulin exchange at MT plus ends is so far little explored.

Here, using linked computational approaches, we analyse the electrostatic fields of TOGs from the TOG polymerase superfamily and simulate their binding trajectories. We show that electrostatic steering accurately guides each TOG domain to a position very close to its crystallographic binding site and that different classes of TOG domain show characteristically different binding rates. We also quantify the stability of the complex of TOGs with curved GTP-tubulin, the steric incompatibility of TOG binding to ‘straight’ GTP-tubulin and the electrostatic and steric matching of each TOG with its cognate tubulin. We predict that incorporation of GTP-tubulin into the growing lattice, with straightening of the tubulin imposed by the lattice, is required to complete the catalytic cycle. Our data show that TOGs are electrostatic adaptors that serve very effectively to enhance the ability of MT plus ends to capture GTP-tubulin and catalyse its polymerisation.

## Results

### TOG domains have a conserved charge-landscape

Electrostatic similarity clustering is a method of identifying the cumulative difference in the spatial distribution of electrostatic potentials for a set of proteins [12]. Cumulative difference in this context refers to the summation of the difference between a parent and a family of electrostatic potentials at each grid point. Comparing the Electrostatic Similarity Index (ESI) distribution for different TOG domain structures show conservation in the electrostatic fields formed by the line of positively charged intra-HEAT loops on each TOG. (**Fig. 2A and B**). The highest ESI scores are located in HEAT repeats (HR) A-D, which engage with β-tubulin. The highest conservation values of all are found in the T1 loop of HR A. In many TOG structures, this is the site of a solvent-exposed tryptophan that when mutated to alanine inhibits tubulin binding [13].

**Fig 2.**
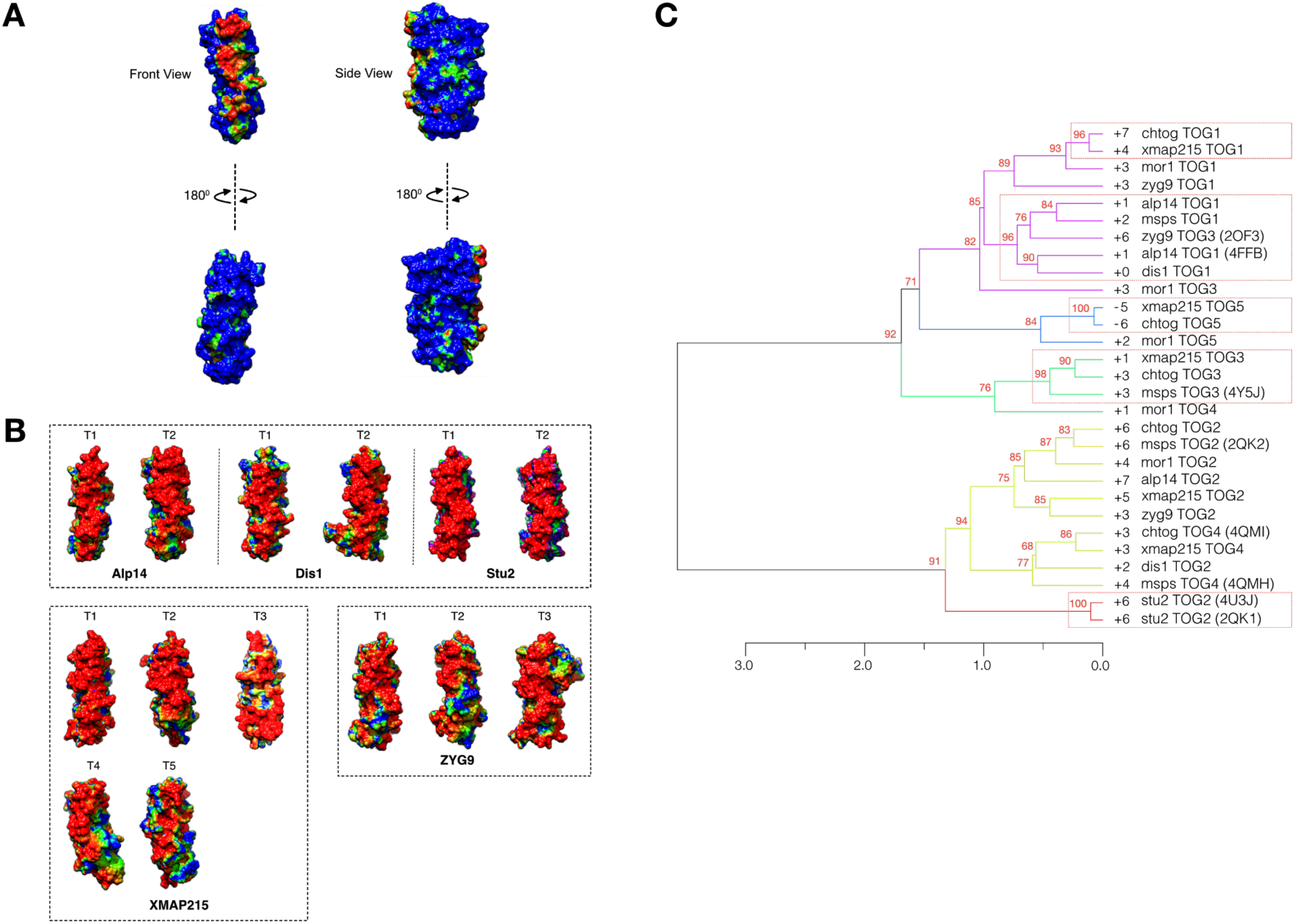
Electrostatic subclasses of individual TOG domains. (A) Cumulative electrostatic similarity distribution, highlighting similarities in electrostatic field strength across all TOG domains in this study. Colour transitions from blue-green-red correspond to ESI values of 0.10 to 0.40 from low to high similarity. (B) Mutation-based perturbation maps for five TOG homologs. Perturbation map colour scheme is blue-green-red for ESI values of 0.5 to 0.9 from low to high similarity. Regions with high ESI values show more evolutionary resistance to computational point mutations of charged residues. (C) Electrostatic clustering analysis by electrostatic similarity distance. Coloured branches in the dendrogram show clusters of similar electrostatic potential. The dashed box represents clusters with p-values greater than 95% confidence by bootstrap resampling.

Comparing the ESI patterns for different TOGs revealed 5 subclasses, partly reflecting the position of each TOG in the primary sequence of its parent polymerase (**Fig 2C**). In the dendrogram, TOG1 domains form a single cluster irrespective of species. TOG2 of Stu2, crystallised alone (2QK1) and in complex with tubulin (4U3J), clusters closely with other TOG2s, except those of Dis1 and Zyg9. TOGs 3,4 and 5 cluster less strongly, but clustering is nonetheless still evident. Electrostatic self-similarity across the TOG protein superfamily is thus strongest for TOGs 1 and 2. This is consistent with experiments on XMAP215 where TOGs 1 and 2 have been shown to make the dominant contribution to the polymerase activity [14], and with recent work contrasting the structure and role of Msps TOG5 with TOG1 and 2 [15].

Primary sequence conservation in TOGs is strongest in the intra-HEAT repeat loops [16, 17], suggesting that conservation of the TOG electrostatics may rely heavily on sequence conservation in these regions. To test this point, we used computational alanine scanning mutagenesis. We found that for all TOGs tested, the field strength at large radii is relatively insensitive to single residue substitutions. This suggests that whilst conservation of the large scale electrostatic field is one driver for sequence conservation, a more important driver is the need to maintain short-range electrostatic interactions with tubulin in the final TOG-tubulin complex.

### Combining PCA and NMA identifies structural-mechanical sub-classes of TOGs

To probe for structural, as opposed to electrostatic, subclasses amongst TOG-polymerase family TOGs, we used principal component analysis (PCA), which quantifies the structural variation amongst TOGs relative to a common invariant core. Figure 3A shows the projection of the structures plotted onto the two largest principal components. This lower dimensional representation groups the structures into four distinct clusters on the basis of the root mean squared displacement (RMSD) of their atom positions. TOG2 and 3 of Msps and TOG2 of Stu2 form a single cluster (blue, green) whereas Zyg9 TOG2 (red) is more closely related to TOG2 and 3 of Msps and TOG2 of Stu2 (green). Thus, 4 distinct, structurally-conserved TOG subclasses are detectable by PCA, suggesting that individual, conserved functional subclasses may also exist.

**Fig 3.**
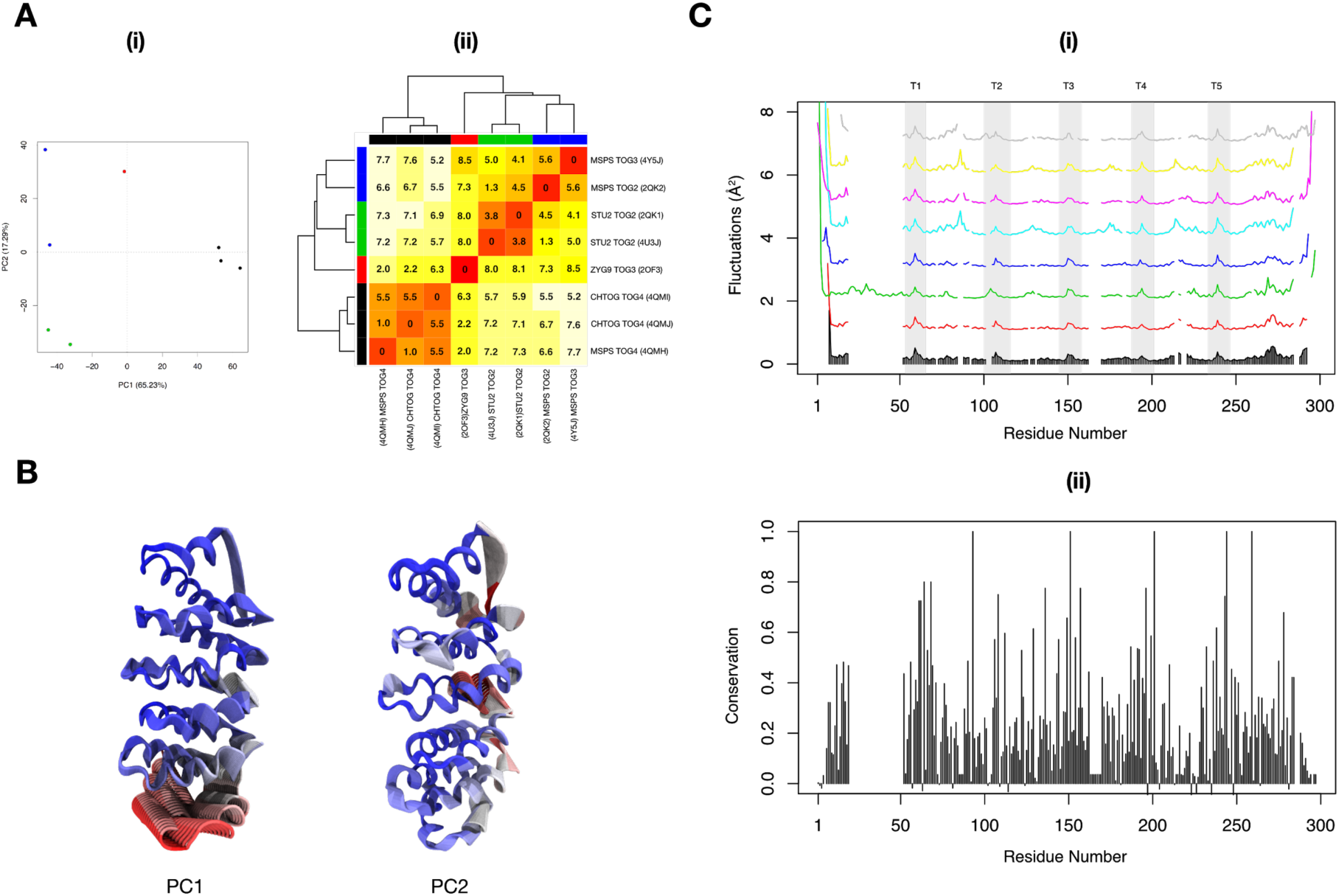
Conservation and variation of structural flexibility identify distinct TOG subclasses. (A) Figure shows the clustering of TOG domains over the first two principal components (i), and a dendrogram and heatmap (ii), both by RMSD. TOG domains form distinct clusters that correlate across species to their position within the TOG array. Zyg9 TOG3 is an outlier, with a RMSD of 8.5 A to its nearest array-positional neighbour Msps TOG3, as a result of its unique N-terminal structure. (B) Visualisation of the first and second principal components of the common structural core of all superimposed structures. The colours depict from red to blue the highest structural deviations between the most dissimilar structures in the set of TOGs. The greatest contributions to each principal component are seen to come from the C-terminal regions of each TOG domain. (C) Atomic fluctuations of the first 200 normal modes for 8 crystal structures (i). All TOG domains show similar fluctuation patterns across species. Sequence conservation from multiple sequence alignment (ii). Most of the cluster-specific variation occurs within the regions that overlap the T1-T5 binding loops. No statistical significance was found between the fluctuation patterns of any one species.

**Fig 4.**
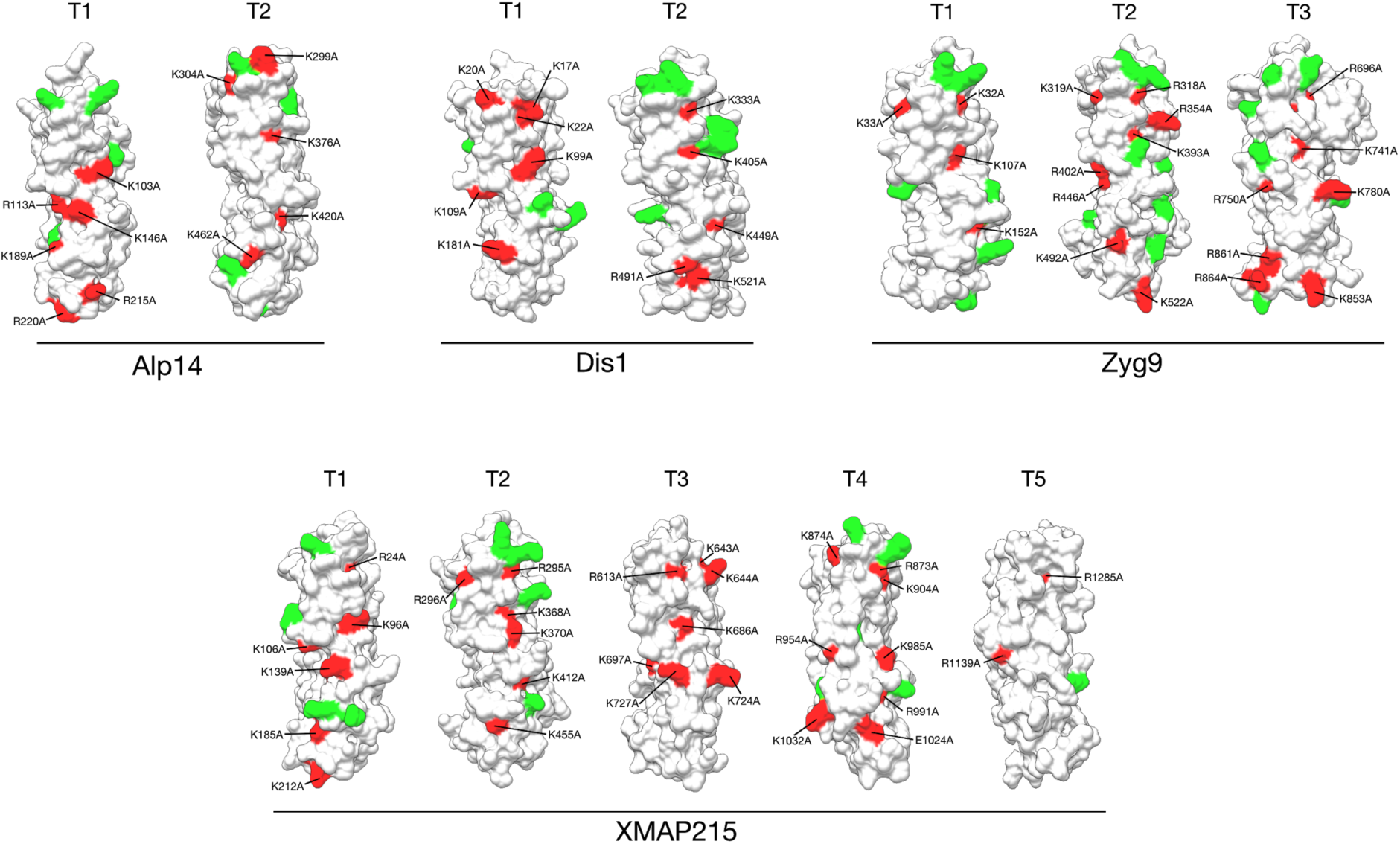
Electrostatic binding free energies of TOG-tubulin complex. Hotspots at which mutation to alanine substantially influences the free energy of binding of each TOG to its cognate tubulin. Red indicates a binding free energy change of ≥ 5 kJ/mol and green ≥ 3 kJ/mol. Residues highlighted in red are indexed relative to the full length protein sequence from the UniProt database. The most significant changes in ΔΔG were found in helical residues surrounding the tubulin binding loops which project outwards from the domain.

Projection of the structures along the first two principal components reveals that, amongst the regions that are common to all TOGs sampled, most of the structural variation between the domains occurs in the C-terminus (**Fig 3B**). The N-terminus remains invariant due to its importance for maintaining tubulin binding and is a likely site for evolutionary modulation of tubulin affinity. This feature is apparent whether or not sampled crystal structures showing an additional secondary structure outside of the TOGs active site. We suggest these helices function to maintain the TOG’s rigid core, especially in the T1-T2 binding loops, rather than endowing the domains with additional functionality.

Aside from their electric fields and their tertiary structures, a third potential determinant of the reactivity of TOG domains is their mechanical stiffness. To assess this, we used normal mode analysis (NMA) to calculate atomic fluctuation profiles of six representative TOG domain proteins (**Fig 3C**). As expected, residues in the tubulin binding loops are all solvent-exposed and mobile, with a higher degree of sequence conservation. The T1 and T3 binding loops, which contain conserved residues that engage with β- and α-tubulin, show slightly larger displacements than the other loops. Crystal structures obtained from isolated TOGs (PDB: 2QK1, 2QK2, 4Y5J) are very similar to those obtained from TOG-tubulin complexes with an RMSD of 0.72 A (see **Supplemental S1**), consistent with the TOG domain acting as a stiff template that can constrain and potentially restrict the conformation of its tubulin binding partner. No direct experimental evidence to date suggests that TOG domains themselves undergo conformational change upon tubulin binding. This, together with our results, further indicates that TOGs have intrinsically low flexibility, across species.

### Mutation of charged residues along the TOG-tubulin interface reduces electrostatic free energies of binding

The different yet conserved electrostatic fields on different TOGs suggested to us that each TOG might be electrostatically matched to a particular tubulin partner. To investigate this point, we calculated electrostatic free energies for the TOG:αβ-tubulin complexes based on the magnitudes and perturbation of these energies when each residue was systematically mutated to alanine. Each structure underwent MD energy minimisation and refinement (see Methods) as a complex. Individual residues were identified that led to a >5 kJ / mol change in the free energy. Positive ΔΔG binding values correspond to a gain in the free energy of association, while negative values correspond to a loss for that residue. Subsequently, residues were ranked and two mutations from each TOG domain were identified as inputs for BD simulations. The most significant changes in binding free energy were found in charged residues in and around the intra-HEAT-repeat loops. This supports the findings of many loss-of-function and binding affinity studies [6,2,18].

### BD shows that electrostatic steering accelerates the capture of tubulins by TOGs

To examine the potential role of electrostatics in long-range guidance of TOGs to tubulin, we used Brownian dynamics. BD trajectories of Stu2 TOG2 diffusing in the electrostatic field of tubulin with and without the presence of the tubulin C-terminal tails were generated at 100 mM monovalent ionic strength to investigate the TOG-tubulin association pathway. Trajectories were generated with no reaction criteria until escape in order to fully sample the electrostatic field (see Methods). Stu2 TOG2 was chosen because of the existence of a high-resolution crystal structure of Stu2 TOG2 bound to tubulin, and extensive biochemical characterization [2, 3]. A random subset of 50,000 trajectories from a total of 1,000,000 was used to prepare the density maps (**Figure 5**).

**Fig 5.**
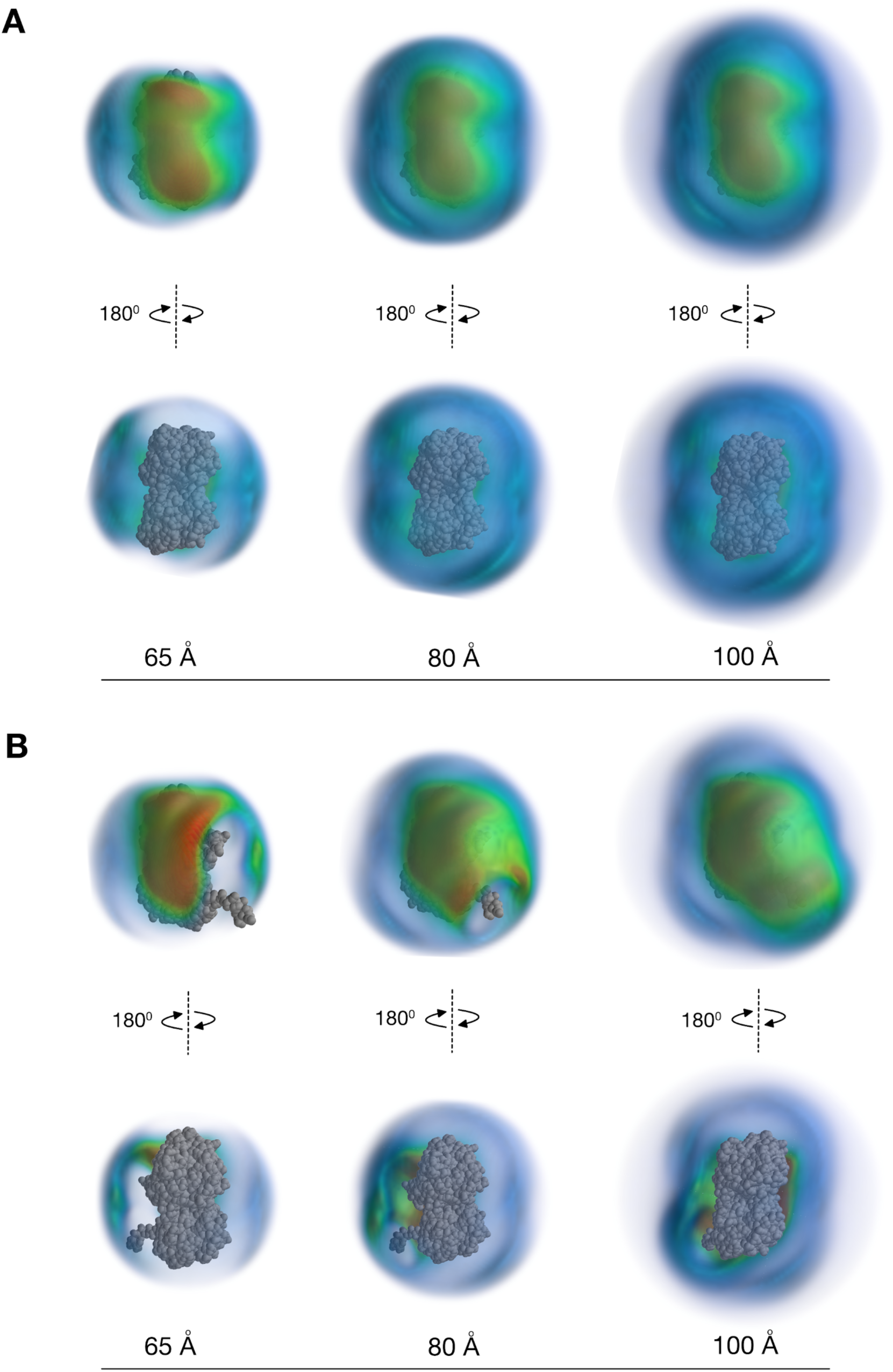
Density maps of TOG diffusion under the influence of a tubulin dimer demonstrate TOG domains undergo strong electrostatic guidance during the diffusional approach to their substrate. The colours represent the average occupancy value for each point in 3D space, with low occupancy in blue and high occupancy in green-to-red. The tubulin dimer is shown at the centre of each plot in grey. **(A)** Figure shows density maps of TOG diffusion under the influence of a tubulin dimer without a C-terminal tail. The TOG domain shows a clear preference for a single binding site that straddles across the intra-dimer interfaces in agreement with the known crystallographic TOG-Tubulin complex (PDB: 4U3J). **(B)** Figure shows density maps of TOG diffusion under the influence of a tubulin dimer with C-terminal tail. Tubulin with C-terminal extensions is shown in the centre of each map in grey. The inclusion of C-termini narrows and focuses the diffusional search made by the TOG domain for its binding site and electrostatic steering begins at higher radius. These maps suggest that the TOG domain undergoes strong electrostatic guidance during the diffusional approach to its substrate. These density maps demonstrate that tubulin C-terminal tails are not required for electrostatic steering but enhance it by expanding the capture radius.

For these studies we used the straight, taxol-stabilised, GDP-tubulin structure (PDB: 1JFF) instead of the tubulin dimer from the TOG-tubulin crystal complex (PDB: 4U3J). The taxol was removed from the structure and energy minimisation and refinement performed (see Methods). This energy-minimised straight tubulin structure was chosen in order to reflect the conformation of GTP-tubulin in free solution, and to avoid biasing the simulation towards formation of the crystallographic complex. As a control we removed the influence of the electrostatic force by increasing the salt concentration and compared this to a model of spatial randomness (**Figure S3**)

Overall, the TOG domain shows a clear preference for a single binding site on our straight tubulin structure (**Figure 5**). This hotspot straddles the α- and β-tubulin monomers. For TOGs binding to tubulin without its C-terminal tails, the hotspot is larger (Fig. 5A) and consequently less dense in comparison to the tubulin with the tails included (Fig. 5B). This implies that at close range, when the TOG domain is close to its final binding position, the inclusion of the tail focuses the TOG domain to a narrower binding site. As the radius of the density maps was increased, the tubulin without tails showed little change in the shape of the binding hotspot, with similar levels of density around the rest of the dimer. In contrast, for the tubulin dimer with tails, the binding hotspot at high radius expanded and began to encapsulate the C-terminal extension of the tubulin, with low density in its surrounding regions. C-terminal tails are thus not required for electrostatically-steered diffusion to capture at the crystallographic binding site, but serve to increase the effective capture radius.

### Translational and orientational degrees of freedom reduce progressively as TOGs approach tubulin

Diffusion of the Stu2 TOG2 domain to tubulin was simulated using BD with formation of the end state defined as the formation of three crystallographic contacts. The translational and rotational coordinates at each time point were computed separately with respect to a spherical coordinate system at set distances from the tubulin active site. For all sampled trajectories, translation of the TOG became constrained at high radius so that the final binding surface faced the tubulin dimer (**Figure 6**). Thereafter, rotational motion was progressively constrained as the radius decreased, and became heavily constrained in the occupancy hotspots immediately overlying the final (crystallographic) site. Rotational restriction tends to occur most strongly when close to the final binding site and helps to rotate the TOG domain into the correct position in the final moments before binding.

**Fig 6.**
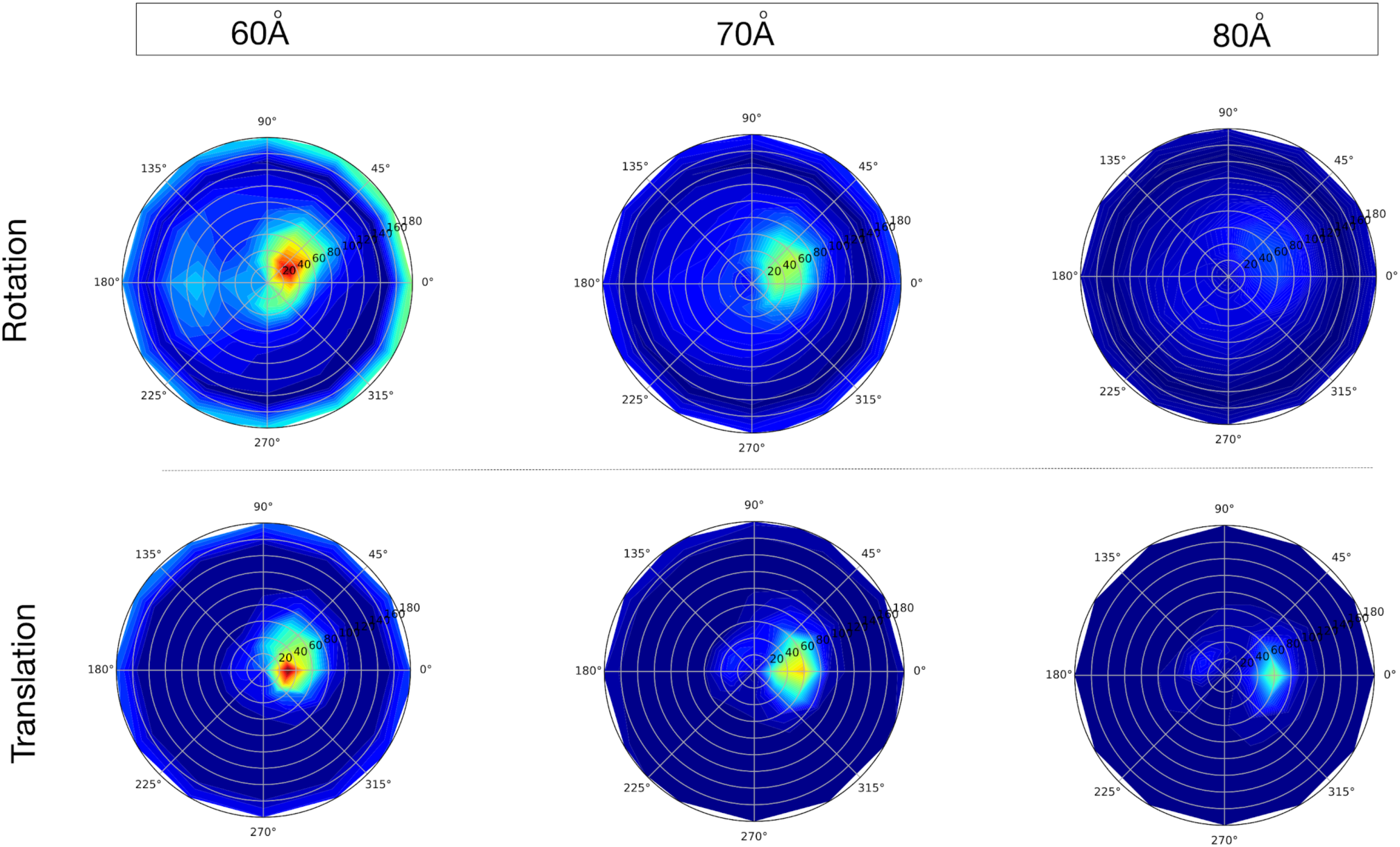
Occupancy maps of associated trajectories as a function of contact distance for translational and rotational coordinates. Red shows high occupancy and blue shows low occupancy. The angles are defined as follows: *θ* is the azimuthal angle in the xy-plane from the x-axis with 0 ⩽ *θ* ⩽ 2$ and % is the zenith angle from the positive z-axis with 0 ⩽ % ⩽ $. Further from the binding site, both rotational and translational coordinates of the TOG domain experience weaker long-range electrostatic forces and lower occupancy values. As the TOG domain approaches its final binding site, substantial electrostatic steering of the TOG domain occurs with translational steering extending over many angstroms whilst rotational restriction tends to occur closer to the final binding site.

### Simulated association rates reflect the differing functional roles for TOG domains

In order to test the relationship between TOG domains and any individual roles of the domains, we calculated ionic strength dependence of the Tubulin-TOG association rates. This allowed us to test whether there was any interspecies difference in kinetics between TOG homologs. We also looked to test whether any differences occurred between individual TOG domains from the same species and whether these differences occurred over an ionic strength range. We performed BD simulations to calculate the association rate of the TOG-tubulin encounter complex.

Figure 7A shows the TOG domain family association rates calculated from these BD simulations. In general, the fastest association rates were determined for TOG1 and TOG2 across species. Multiple biochemical studies have established the functional equivalence of TOG1 and TOG2 for tubulin binding and polymerase activity [3,14,15,19]. Interestingly, for the TOG isoforms in *S. pombe* cells, Dis1 TOG1 associates more slowly than Dis1 TOG2. The differences in the simulated association rates are consistent with the differing functional roles of each protein in cells. Alp14 is closely associated with enhancing MT growth during bipolar spindle formation. Dis1 is known to promote MT bundling alongside the Ndc80 kinetochore protein. Zyg9 TOG3 was unusual in having a higher association rate to *C. elegans* tubulin than Zyg9 TOG1 and Zyg9 TOG2. Zyg9 TOG3 consists of an additional N-terminal heat repeat that binds laterally across the domain and may account for the simulated differences in rate. XMAP215 TOG1 was found to have the highest simulated association of all species under consideration. XMAP215 TOG1 has a large overall negative charge which promotes strong electrostatic steering. Overall this investigation suggests that the higher association rate constants for TOGs 1 and 2 reflect their known preference for binding to unpolymerised tubulin.

**Figure 7.**
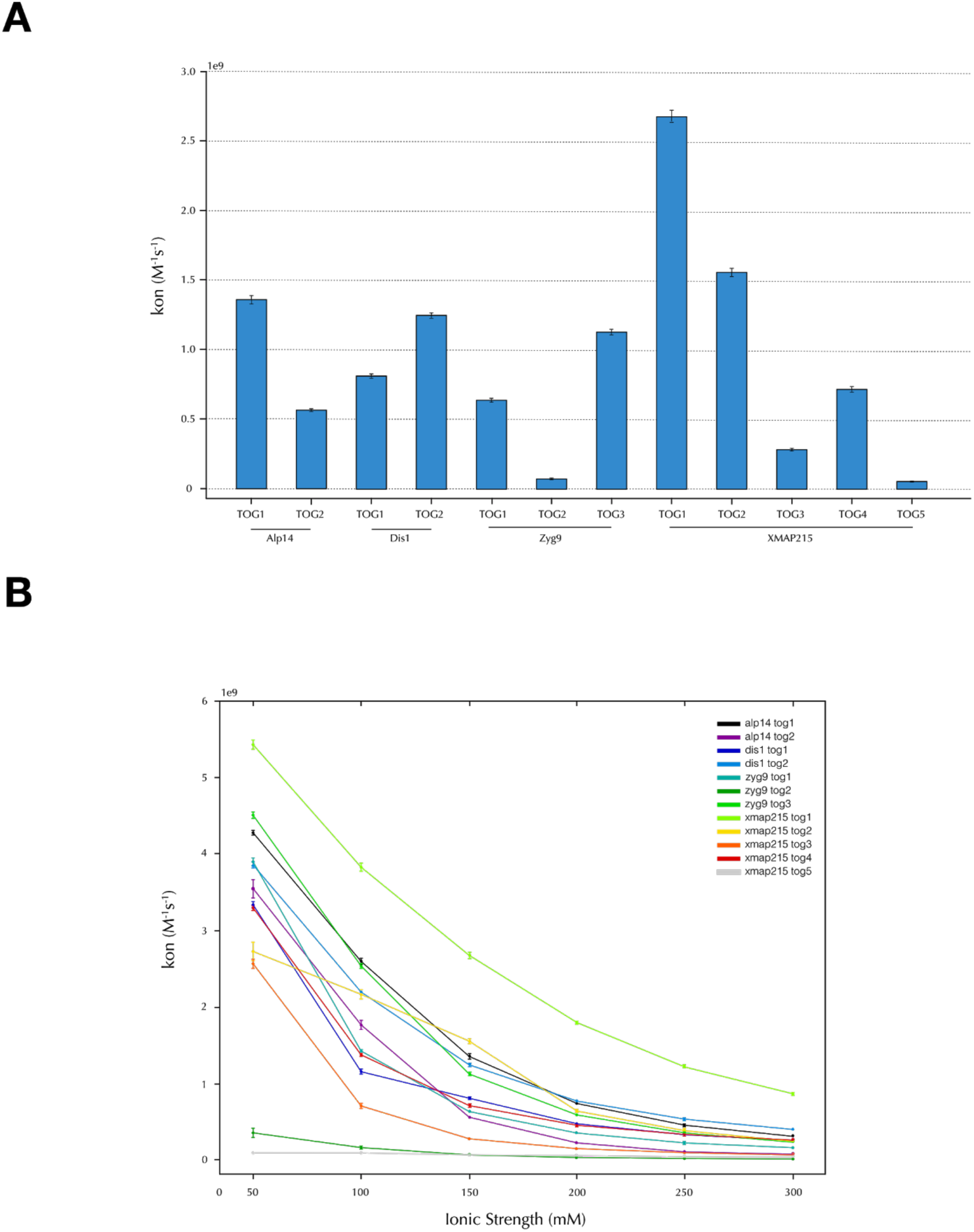
BD simulations reveal inter- and intra-species variability in TOG-tubulin association rates. (A) TOG association rates for each TOG family member at 150 mM ionic strength. We saw both inter- and intra-species variability in the association rates. In general, the fastest association rates were determined for TOG1 and TOG2 across species. Error bars in both plots depict standard error of the mean. (B) To test if these differences were consistent across an ionic strength range, we calculated the association rate of TOG to tubulin at a concentration of between 50-–300 mM monovalent salt. Association rate constants for all TOG domain species to their cognate tubulin reduce as the ionic strength increases.

### Electrostatic field strength of the TOG domain is a key determinant of its association with tubulin

From the alanine mutagenesis performed in Figure 4, binding to tubulin of two high-scoring mutant TOG structures was simulated. We found that mutations that change the electrostatic binding free energy to the final complex with curved tubulin also strongly affect the rate constants for the diffusion to capture reaction, showing that the strength of the electrostatic attraction is the dominant determinant of reaction rates.

### Protein-protein docking identifies an energetically favourable binding position that correlates with the crystallographic position

The near-native TOG-tubulin complex was predicted using rigid body docking, using only a simple scoring function based upon electrostatics and desolvation. This result further supports our hypothesis that electrostatics are a critical/key determinant in TOG-Tubulin binding kinetics as the near native final complex formed in absence of further short range interactions. Table 2 shows a clear correlation between the interface RMSD and the energy score. This relationship holds within species and is TOG array position dependent. For instance, the correlation coefficient identifies the difference in association rate between Alp14 and Dis1, and the electrostatic dissimilarity of XMAP215 TOG5.

**Table 1.** Charged-to-alanine computational point mutations on calculated TOG domain association rates reduced the association rate for all domains. Two mutations of charged residues for each TOG domain were selected with a cutoff of > 5 kJ/mol. Association rates are presented with 95% confidence intervals. Predominantly, each mutation had the effect of significantly reducing the association rate for all domains.

| Domain | Residue | BD $k_a$ | WT BD $k_a$ | ΔBD $k_a$ |
| --- | --- | --- | --- | --- |
| Alp14 TOG1 | K146A | $5.5e+8 \pm 0.15$ | $1.36e+9 \pm 0.03$ | $-8.10E+08 \pm 0.153$ |
| | R215A | $3.8e+8 \pm 0.2$ | | $-9.80E+08 \pm 0.202$ |
| Alp14 TOG2 | R376A | $9.0e+7 \pm 0.7$ | $5.65e+8 \pm 0.17$ | $-4.75E+08 \pm 0.720$ |
| | R462A | $2.4e+8 \pm 0.1$ | | $-3.25E+08 \pm 0.197$ |
| Dis1 TOG1 | K17A | $2.1e+8 \pm 0.1$ | $8.06e+8 \pm 0.16$ | $-5.96E+08 \pm 0.189$ |
| | R22A | $1.23e+8 \pm 0.1$ | | $-6.82E+08 \pm 0.189$ |
| Dis1 TOG2 | K449A | $9.1e+8 \pm 0.2$ | $1.25e+9 \pm 0.02$ | $-3.40E+08 \pm 0.200$ |
| | R491A | $9.0e+8 \pm 0.2$ | | $-3.50E+08 \pm 0.201$ |
| Zyg9 TOG1 | R32A | $1.1e+8 \pm 0.07$ | $6.40e+8 \pm 0.14$ | $-5.30E+08 \pm 0.157$ |
| | K33A | $5.5e+7 \pm 0.5$ | | $-5.85E+08 \pm 0.519$ |
| Zyg9 TOG2 | R318A | 0 | $7.39e+7 \pm 0.63$ | $-7.39E+07 \pm 0.630$ |
| | K338A | $3.64e+8 \pm 1.11$ | | $2.90E+08 \pm 1.276$ |
| Zyg9 TOG3 | K741A | $4.1e+8 \pm 0.2$ | $1.13e+9 \pm 0.02$ | $-7.20E+08 \pm 0.201$ |
| | K853A | $6.13e+8 \pm 0.3$ | | $-5.17E+08 \pm 0.301$ |
| XMAP215 TOG1 | K96A | $2.28e+9 \pm 0.03$ | $2.68e+9 \pm 0.04$ | $-4.00E+08 \pm 0.050$ |
| | K139A | $2.33e+9 \pm 0.03$ | | $-3.50E+08 \pm 0.050$ |
| XMAP215 TOG2 | K370A | $1.5e+8 \pm 0.09$ | $1.16e+9 \pm 0.03$ | $-1.01E+09 \pm 0.095$ |
|  | R455A | 7.9e+8 ± 0.2 |  | -3.70E+08 ± 0.202 |
| XMAP215 TOG3 | R613A | 7.6e+6 ± 1.6 | 2.85e+8 ± 0.14 | -2.77E+08 ± 1.606 |
|  | K727A | 3.21e+7 ± 0.3 |  | -2.53E+08 ± 0.331 |
| XMAP215 TOG4 | R954A | 2.6e+8 ± 0.09 | 7.18e+8 ± 0.21 | -4.58E+08 ± 0.228 |
|  | E1024A | 8.6e+8 ± 0.2 |  | 1.42E+08 ± 0.290 |
| XMAP215 TOG5 | R1285A | 2.0e+7 ± 0.2 | 5.50e+7 ± 0.43 | -3.50E+07 ± 0.474 |
|  | R1339A | 2.7e+7 ± 0.3 |  | -2.80E+07 ± 0.524 |

**Table 2.** Prediction of the TOG-tubulin complex by rigid body docking. PyDock results show correlation between interfacial root mean squared displacement (iRMSD) and energy score. iRMSD is the difference between the final docked orientation of the TOG domain and the known TOG-tubulin native complex. The docking results show a clear correlation between the Spearman rank coefficient and on-rate constant calculated from BD simulations. These differences reflect the tubulin-binding properties of individual TOG domains. PyDock shows that TOG domains can bind tubulin in a near-native conformation, without defining a binding criteria. This further supports the hypothesis that electrostatics are the critical determinants of TOG domain binding kinetics.

| Domain | Spearman Rank | p-value | BD k<sub>a</sub> |
| --- | --- | --- | --- |
| Alp14 TOG1 | 0.42 | 1.80e-5 | 1.02e+8 |
| Alp14 TOG2 | 0.20 | 0.043 | 2.51e+7 |
| Dis1 TOG1 | 0.17 | 0.083 | 9.43e+06 |
| Dis1 TOG2 | 0.48 | 4.70e-7 | 2.48e+07 |
| Zyg9 TOG1 | 0.66 | 1.10e-13 | 1.78e+08 |
| Zyg9 TOG2 | 0.36 | 0.00023 | 2.83e+07 |
| Zyg9 TOG3 | 0.50 | 1.50e-7 | 2.36e+07 |
| XMAP215 TOG1 | 0.45 | 2.60e-06 | 2.36e+07 |
| XMAP215 TOG2 | 0.17 | 0.099 | 4.72e+08 |
| XMAP215 TOG3 | 0.37 | 0.0017 | 2.51e+07 |
| XMAP215 TOG4 | 0.35 | 0.00041 | 4.62e+06 |
| XMAP215 TOG5 | 0.29 | 0.003 | 4.30e+06 |

### TransComp Methods confirm that TOGs are electrostatically driven to tubulin dimers

As a control to guard against model-dependent bias of our electrostatic steering simulations, we used transient complex theory [20] to make a model-independent test for electrostatic guidance. The calculated association rates (**Fig. 9**) show similar patterns of conservation to the prior BD simulations, with the highest association rates found for TOG1-2, confirming that the TOG-tubulin association reaction is electrostatically driven.

**Figure 9.**
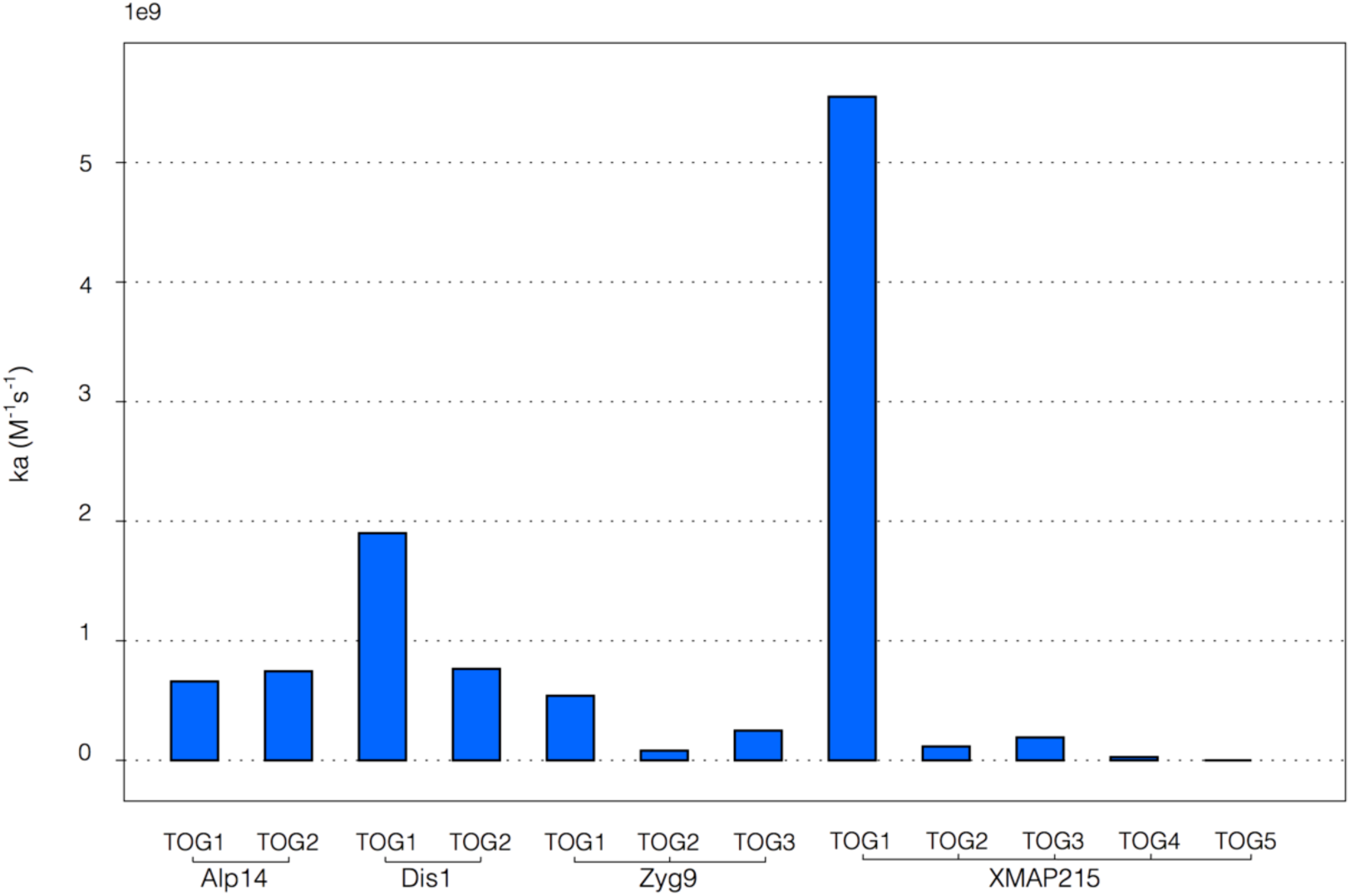
Predicted association rates for TOG domains derived from transient complex theory. For associating proteins the transient complex is defined as the point when both proteins have near-native separation and relative orientation but are yet to form short range interactions. TransComp shows comparable results to the association rates calculated from BD simulations.

## Discussion

TOG family MT polymerases accelerate the net GTP-tubulin on-rate at MT plus ends by up to 10-fold [1]. TOGs bind tightly to tubulins from diverse sources - for example, *S. cerevisae* TOGs have been used for affinity chromatography of tubulins from multiple species [10]. However for the TOG-driven acceleration of MT plus end growth to be effective, TOGs need not just to bind rapidly to tubulin, but also to dissociate rapidly. There is good evidence that individual TOG-polymerase molecules are processive, tracking MT plus ends as they grow, repeatedly recruiting tubulin heterodimers from solution and inserting them into the growing tip-lattice. The maximum rate of TOG-catalysed plus end growth of MTs at full TOG-polymerase occupancy is at least 50 nm s^-1^, and might be considerably more [19]. But taking this value, and assuming one TOG-polymerase per protofilament, each TOG polymerase is assembling ∼7 GTP-tubulin heterodimers per second per protofilament, which implies that the entire tubulin capture-and-assemble cycle time is complete on average within <140 ms.

TOG-protein catalysis of MT plus end growth appears most effective for matched pairs of TOGs and tubulins from the same organism [15, 21]. The best studied TOG-polymerase, *Xenopus* XMAP215, behaves as a classical enzyme that accelerates the reaction that it catalyses (GTP-tubulin exchange) but does not alter its equilibrium constant [1]. By contrast *S. pombe* Alp14, a TOG family polymerase from *S. pombe*, both accelerates exchange and shifts the exchange equilibrium in favour of polymerisation [21]. In common between these two types of mechanism is the imperative not only to bind GTP-tubulin from solution as rapidly as possible, and then also to donate it efficiently into the growing tip lattice. How does this work?

Our structural and electrostatic analyses of TOGs from TOG family MT polymerases reveal them to be stiff modules whose electric fields have a distinct structure that is characteristic of each TOG subclass. We initially envisaged potentially two stages of TOG-tubulin binding, both involving the TOG electric field – we postulated longer-range steering / guidance interactions that attract tubulin into the TOG active site, followed by short-range ionic interactions with the tubulin substrate. In the event we identified 3 distinct components of the TOG-tubulin interaction mechanism, firstly long range electrostatic attraction (enhanced by the tubulin E-hooks), secondly a short range residue-specific electrostatic match of the TOG to its cognate tubulin, and finally conformational selection, made possible by the stiffness of the TOG as a binding partner, whereby GTP-tubulin needs to bend in order to make the final crystallographic complex with the TOG.

This last conclusion derives from coarse-grained NMA, which indicates that individual TOG domains are relatively rigid. This gives confidence that our BD simulations, which treat the diffusing molecules as rigid bodies, are reasonably realistic. It also suggests that TOGs can act as engines for conformational selection, in that the stiffness of the TOG imposes a requirement that GTP-tubulin must adopt a curved conformation in order to make the tightest possible complex, with all electrostatic contacts satisfied, as in the crystallographic complex.

Our work shows that for straight tubulin heterodimers, the long-range electrostatic steering process and the shorter range residue-specific electrostatic match of the TOG to ‘straight’ tubulin are both very similar to their equivalent processes with curved tubulin. However the third class of interactions, the close-range ‘fit’ of the TOG protein to tubulin, is very different. As discussed by Rice and Brouhard [22] and by Byrnes and Slep [15], the binding of TOGs to straight tubulin leaves an unfilled cavity. Our finding that TOGs are stiff means that TOGs can potentially act as rigid templates to drive incoming GTP-tubulin subunits into a curved conformation by conformational selection. In this way the the TOG-tubulin binding interface can be extended, and the stability of the complex maximised. Our electrostatic free energy calculations lend further weight to the view that this final complex is highly stable.

### Proposed catalytic mechanism of TOG-family MT polymerases

The rapid, electrostatically-steered recruitment of GTP-tubulin to TOG domains, and the stability of the resulting TOG-tubulin complex together beg the question, how is GTP-tubulin released from the TOG so that it can assemble into the growing MT? Our work has not directly addressed the mechanism by which TOGs donate tubulin into the MT tip-lattice, but given the stability of the final TOG-tubulin complex, we infer that a competitive process must be introduced to drive release of the tubulin and recycling of the TOG for another round of catalysis. We propose that this release mechanism is a matched conformational selection process imposed by the growing MT tip-lattice itself. **Figure 10** shows a schematic of this mechanism.

**Fig 10.**
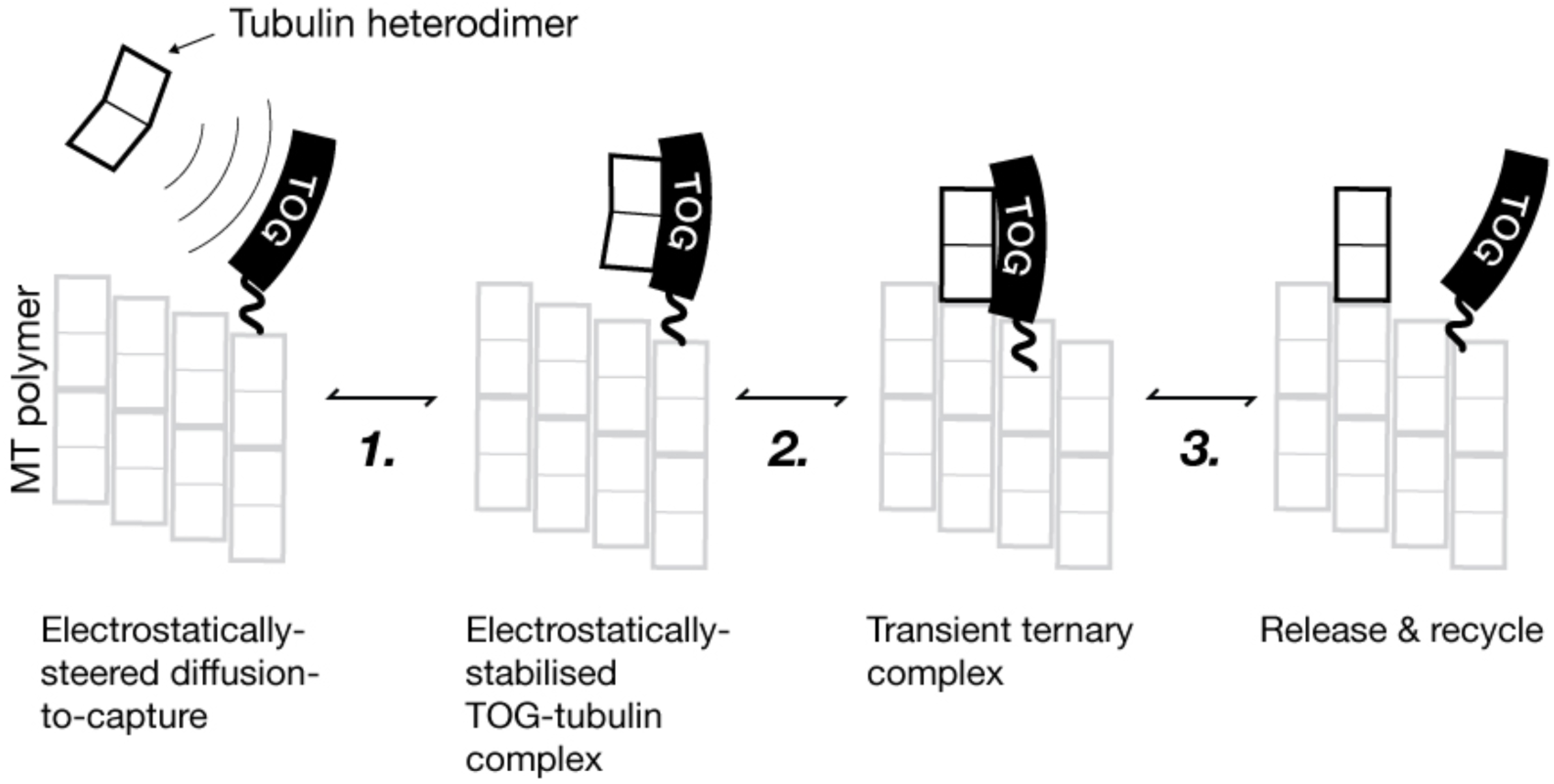
Proposed 3 step scheme for TOG-catalysis of MT plus end growth. We envisage that following electrostatic steered diffusion to capture (**1**), incoming GTP-tubulin heterodimers undergo conformational selection driven by stereospecific binding to the more rigid TOG domain, effectively locking them into a conformation that favours their binding into unoccupied sites in the MT (**2**). The stability of this TOG-tubulin leads us to propose that binding of the TOG-bound GTP-tubulin to its prospective binding site in the tip lattice is required to compete the tubulin away from the TOG (**3**).

Our scheme is minimal in that it does not propose specific roles for the different TOG subclasses of particular TOG-family polymerases (see for example [3]), but it does have an appealing symmetry, in that the zippering of the tubulin to the TOG competes with the zippering of the tubulin into the MT tip. In this way both the on- and off-rates for GTP-tubulin at the MT tip are accelerated by the TOG-polymerase, without much affecting the equilibrium dissociation constant for GTP-tubulin at the plus end tip. Note that *S. pombe* Alp14, which does shift this equilibrium, is a special case in which, as we previously argued [21], a non-TOG C-terminal domain stabilises the polymer and reduces the off-rate for GTP-tubulin. Recent work shows that Stu2, the *S. cerevisae* homologue of Alp14, acts as a MT destabiliser *in vivo* in budding yeast spindles [23], emphasising that TOG-family polymerases accelerate tubulin loss as well as tubulin addition at MT plus ends.

In conclusion, our work establishes that TOG domain MT polymerases work first by providing an optimized landing platform for tubulin, attracting and orienting the incoming GTP-tubulin, and second by shaping and rapidly donating it into the tip lattice at rates that exceed the uncatalysed background rate of GTP-tubulin incorporation into the tip lattice by about an order of magnitude.

## Materials and Methods

### Structure Preparation

TOG domain structures were acquired from the Protein Data Bank (PDB). Missing loop regions and homology models were created and refined using MODELLER (version 9.16) [24]. Template structures for comparative modelling were selected based on closest sequence identity to the target sequence from UniProt. (see Table 3). Up to 50 models were generated per target sequence and ranked by DOPE-HR scoring. The final structures underwent MD refinement and minimisation. All models were validated by PROCHECK to ensure stereochemical correctness of the final structures (see Supplementary Table 1).

### Electrostatic Calculations

Electrostatic potentials were calculated with APBS (version 1.4.1) [25], using AMBER charges and radii at pH 6.9 obtained from PDB2PQR (version 1.9.1) [26]. The full, linear Poisson-Boltzmann (PB) equation was solved for each structure. Atomic charges were mapped to grid points via B-split discretisation. The protein boundary was delineated by a dielectric boundary between solute (ep of 2) and solve (es of 78.54). The surface was defined as the molecular surface (srfm: smol, srad: 1.40). Potential values were calculated on a grid that provided a minimum resolution of not greater than 0.5A.

### Electrostatic Similarity Analysis

Electrostatic potentials for each TOG domain were compared for cumulative similarity and resistance to alanine scanning mutagenesis by the analysis of electrostatic similarity of proteins frameworks [12]. AESOP aligns a set of electrostatic potentials onto a single TOG domain and calculates its cumulative distribution of electrostatic similarity by the following expression:

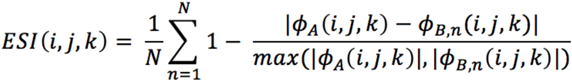

Here ɸ_A_ represents the potential map to which all other potentials are compared, and ɸ_N_ denotes the Nth potential within the set. TOG homologs were compared to the electrostatic potential of 4U3J TOG2. The electrostatic potential of Surface projections of electrostatic similarity were output to OpenDX files and rendered using UCSF chimera (version 1.10.1) [27].

### Normal Mode Analysis

Normal mode analyses were performed using the Bio3d package (version 2.2-2) [28] for each TOG domain. The lowest frequency modes were calculated using a coarse-grained elastic network model (ENM). The residue specific force-field (sdenm) was selected to model the harmonic potential between atoms. For fluctuation analysis each structure was superimposed onto an invariant core and unaligned atoms were retained in the final output.

### Principal Component Analysis

Structures were subject to iterative rounds of structural superposition to identify an invariant core. This region was used as a reference frame for the alignment of all crystal structures and used to calculate pairwise RMSD values. TOG domains were then projected onto a subspace defined by principal components with the largest eigenvalues and labelled into groups by hierarchical clustering. PCA conformers were visually represented as a PBD trajectory. All processing steps were undertaken in R using the Bio3D library (version: 2.2-2) and 3D structures generated with Pymol (version: 1.8.0) [29].

### Electrostatic Free Energy Calculations

Theoretical alanine scanning mutagenesis was performed for all charged residues in each TOG domain. Electrostatic free energies of association were calculated for each TOG mutation, according to the following thermodynamic cycle that includes association in both explicit solvent and vacuum reference states:

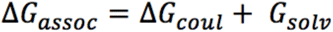

The electrostatic free energy of mutants are represented relative to the parent protein as described:

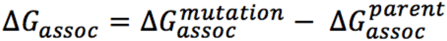

All steps in this cycle were automated through the use of Python scripts using the pandas library (version 0.18.0) [30] and binding plots were generated using USCF Chimera. Side-chains were truncated to alanine using modeller (version 9.16).

### Brownian Dynamics

BrownDye (version: 31 Jan 2016) [31] was used to compute TOG domain associate rates to the tubulin dimer. Atomic charges and radii were generated from PDB structures by pdb2pqr, using AMBER force field at pH 6.9. APBS was used to calculate electrostatic grids by solving the linear Poisson-Boltzmann equation. The reaction criteria were determined from a pilot study of Stu2 TOG1-2:-tubulin complex after MD refinement. Reactions were considered successful when simulations ultimately formed at least three pairwise interactions. Each TOG domain was simulated 200-500K times by the NAM algorithm.

### Molecular Dynamics

Homology models for each component of the TOG-Tubulin complex underwent energy minimisation and equilibration using Amber11 [32]. Missing hydrogen atoms were replaced by the LEaP module and ligand parameters for GTP and Mg were obtained from the Amber parameters database [33, 34]. The system was neutralised by Na^+^ counterions and solvated with TIP3P waters inside of an 12 Å octahedral box. All simulations were run with the Sander module of Amber11, using an ff99SB all-atom force field and obtained parameters for guanine nucleotides and magnesium ligands. Energy minimisation was carried out with decreasing constraints on the structures, followed by constant volume heating to 300 K for ∼10 ps and 200 ps constant temperature and pressure equilibrium. The SHAKE algorithm was used to constrain all hydrogen-heavy bond lengths. Short-range non-bonded interactions were truncated at 10 Å with the Particle-Ewald method.

### TransComp

Rate constants were calculated by a local version of the TransComp method (http://pipe.sc.fsu.edu/transcomp/) [20], according to:

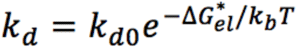

where, k_d0_ is the basal rate for a protein under no influence of long-range electrostatic interactions and G*_el_ is the electrostatic interaction energy normalised by the Boltzmann factor calculated on the edge of the transient complex.

### Binding Maps

Coordinates for each BD trajectory were extracted using the auxiliary program process_trajectories and stored in HDF5 format using PyTables. Density maps were constructed by kernel density estimation from the SciPy package and volume rendered in 3D using Mayavi (version: 4.5.0) [35].

### PyDock

Protein-protein docking of TOG domains to Tubulin dimers was undertaken using PyDock (version: 3.0) [36]. The top 100 structures were ranked by their energy score from 10,000 independent runs. To improve docking results, a single residue restraint was applied to a conserved tryptophan in the first N-terminal helical region of each TOG domain. The RMSD function of USCF chimera was applied between the interfacial atoms of each TOG domains final posed structure and the equivalent atoms in the known TOG:tubulin complex (PDB: 4U3J). The resulting scatter plots and distributions were produced in Python using the Seaborn package (version: 0.7.1) [37].

### Occupancy Maps

The orientational and positional coordinates of each TOG domain relative to the tubulin dimer were computed separately with respect to a reference coordinate system (see Figure S5). For positional coordinates a spherical coordinate frame was chosen. The x- and y-axis are orthogonal to the z-axis. The angle between the z-axis and the center-to-center vector for a given trajectory position is denoted as the zenith angle (**&**). The azimuthal angle (’) is the angle is the angle in the xy-plane from the x-axis. The orientational coordinates use the same spherical coordinates frame but exchange the center-to-center vectors with normal vector to a plane (denoted n1). XYZ trajectory files were processed in Python using the MDAnalysis [38] and NumPy packages [39].

### Tubulin C-termini Modeling

Coordinates from the C-terminal tails of tubulin were kindly gifted by Jack Tuszynski, University of Alberta. These tail coordinates, obtained from the end points of MD simulations [40] were docked to the tubulin structure and converted to an electron density map using UCSF Chimera. New tail sequences for the homology tubulin sequences were generating using Modeller and refined into the density map using MD flexible fitting using DireX (version 0.6.2) [41].

### Calculation of Ripley’s K-function

Ripley’s K function was used to detect the degree of deviation from complete spatial randomness for both physiological and high salt conditions. The function was calculated using the Spatspat package in R [42] and is defined as:

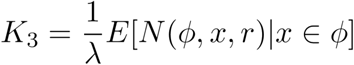

where (is the expected number of points per unit volume and and N(%, x, r) is the number of points of the stationary point process that fall within a distance r of x. The K-function was plotted at multiple distances from a random sample of 5000 trajectories to show how the point distribution changed with scale. The K function of these values, with isotropic edge correction, was plotted. The envelope command was used to compare each 3D point pattern against complete spatial randomness with 99 simulations used to compute the 95% significance level.

## Supporting Information

**Figure S1.**
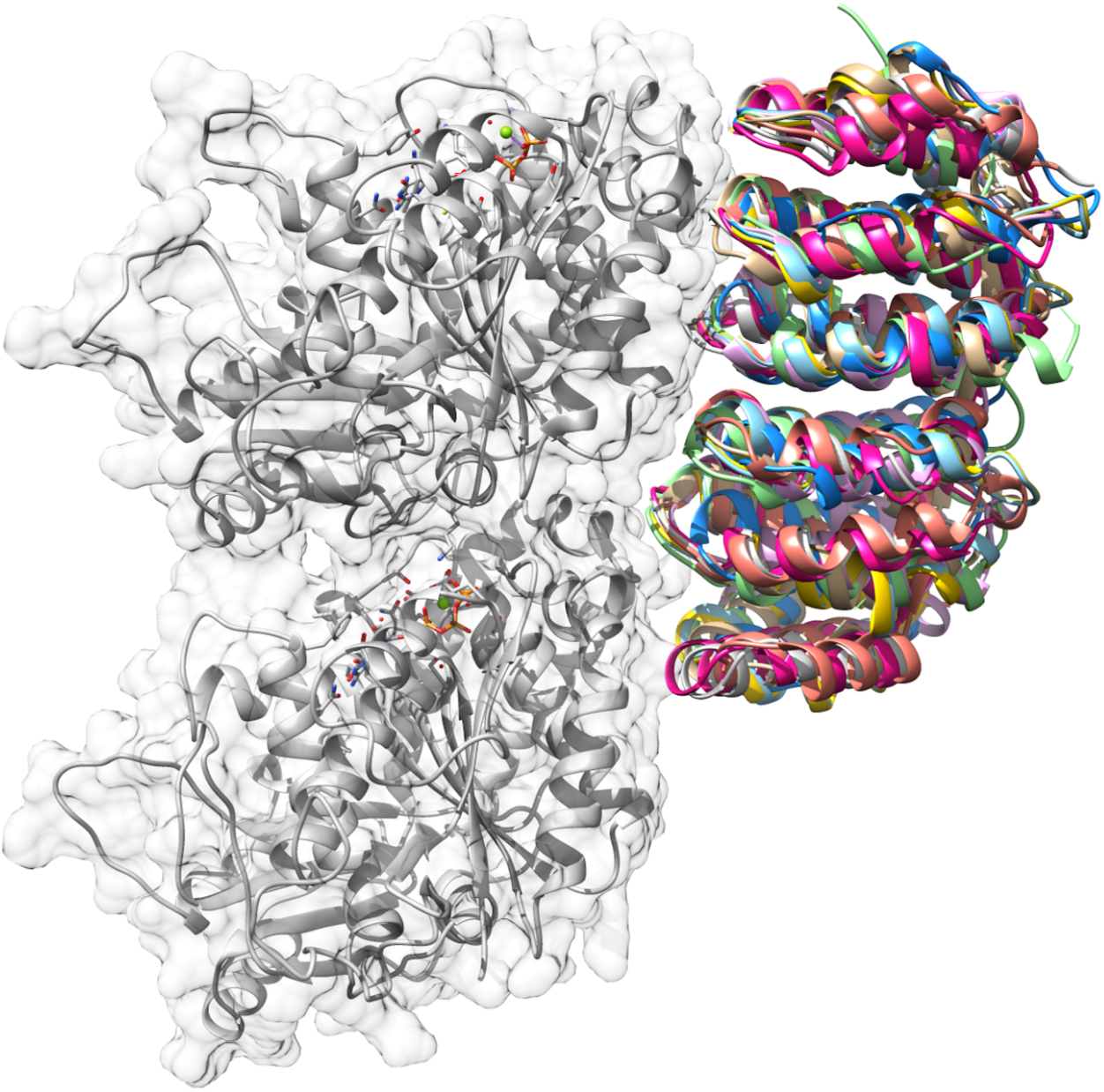
TOG structural alignment of nine crystal structures from XMAP215 family members. Overall the structures aligned with a RMSD of 1.2 Å showing a high degree of structural conservation, irrespective of species and TOG array position.

**Figure S2.** Movie shows 360 degree rotation of density maps of TOG diffusion under the influence of a tubulin dimer, with and without C-terminal tail, at a distance of 80 Å. NoTail Tail

**Figure S3.**
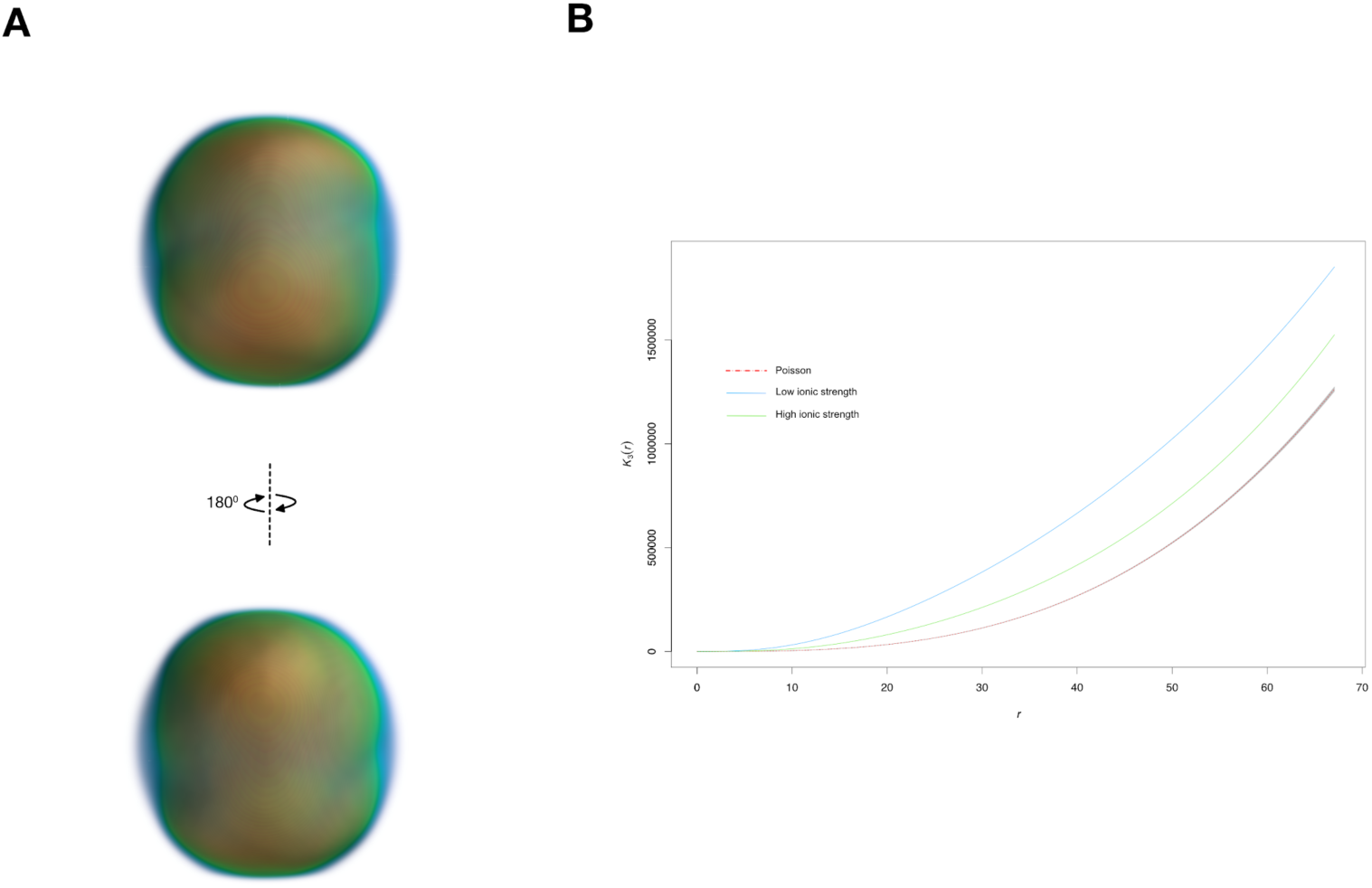
A control of TOG domain diffusion tested against a model of spatial randomness. (**A**) Density map was plotted for a TOG domain diffusing under high salt conditions. The volume representation shows a homogeneous diffusion pattern indicative of Brownian motion. This contrasts with the physiological salt condition, as shown in Figure XX. (**B**) Statistical quantification of TOG domain diffusion under normal and high-salt conditions using Ripleys K-function. The red dashed line shows the expected K-value at the corresponding distance for a point pattern under complete spatial randomness. The green and blue lines represent high salt conditions and physiological salt conditions, respectively. Both of the TOG diffusion regimes show a distinct difference in their overall point distribution for each salt condition with no overlap in their confidence intervals.

**S4.**
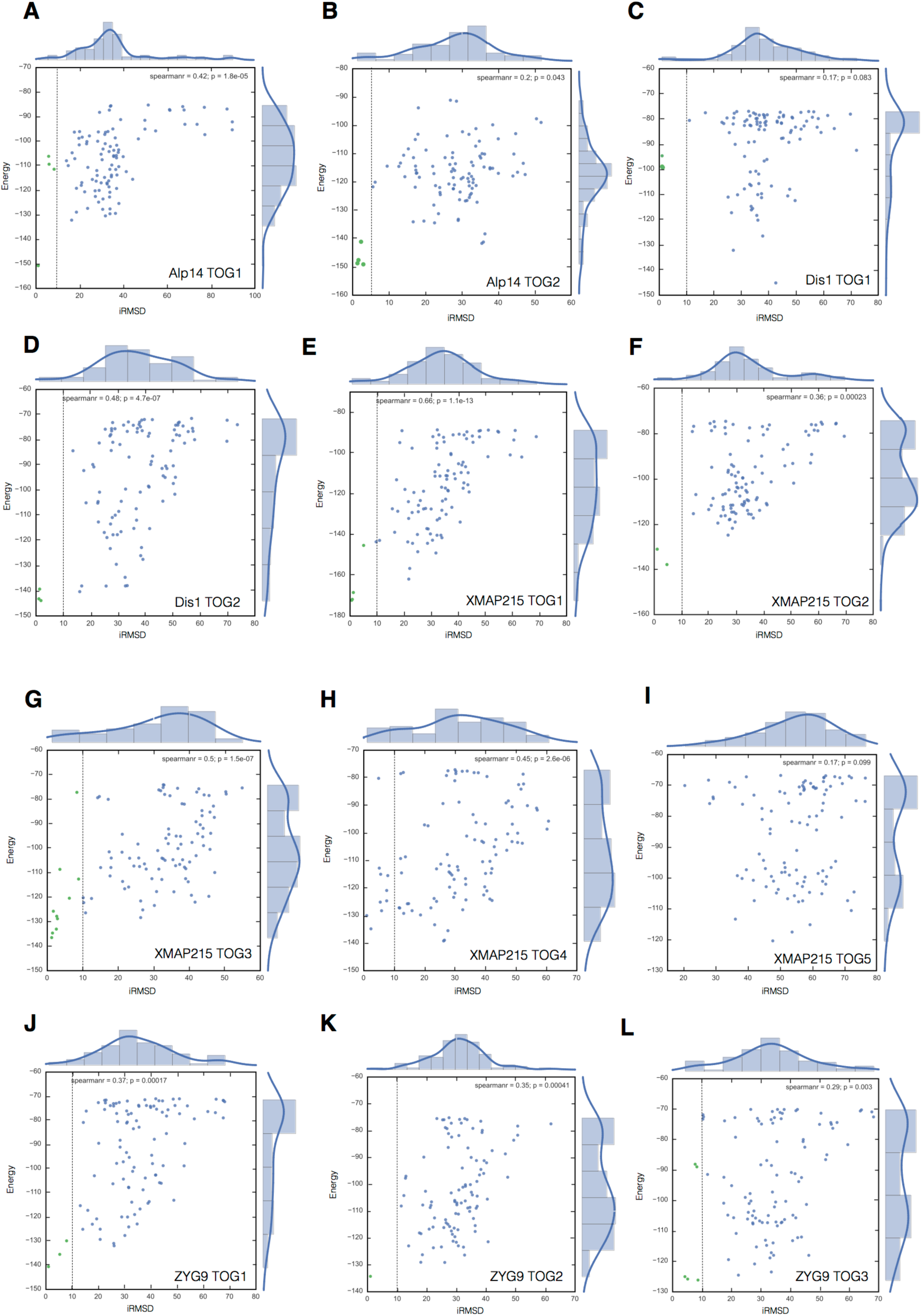
The top 100 docked TOG domains for each species plotted as PyDock energy scores vs iRMSD. Each structure with an iRMSD of below 10 A is coloured green whilst the remaining structures are coloured blue. The plots show a clear relationship between Spearman rank coefficient and energy score, which correlate to those domains with higher simulated association rates.

**S5.**
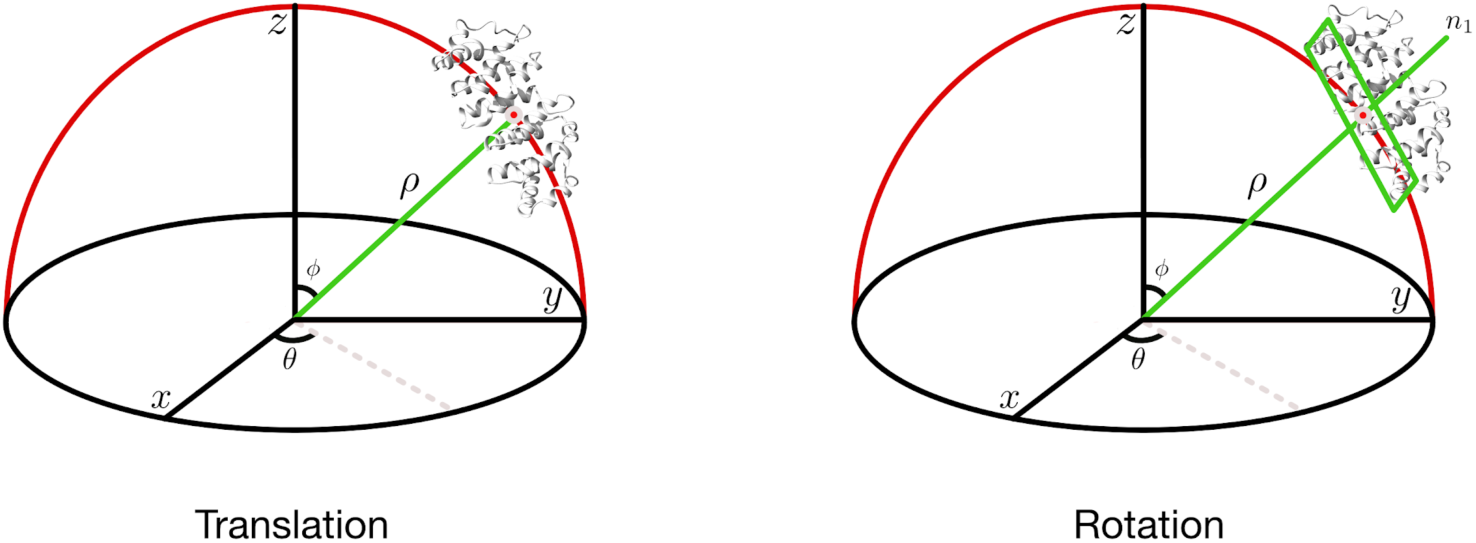
Graphical representation of how the translational and rotational reference schemes were calculated for each BD trajectory. (**A**) For translational coordinates, polar coordinates were determined for each time point based on the domain’s centre-of-mass. (**B**) For rotational coordinates, a plane was fit through the domain’s interfacial atoms and the normal vector to this plane was used to calculated the polar angles for each time point. **<u>S5. These supplementary movies show the binding of a TOG domain to unbond (1JFF) and lattice-incorporated (4U3J) tubulin dimers.</u>**

**T1.**
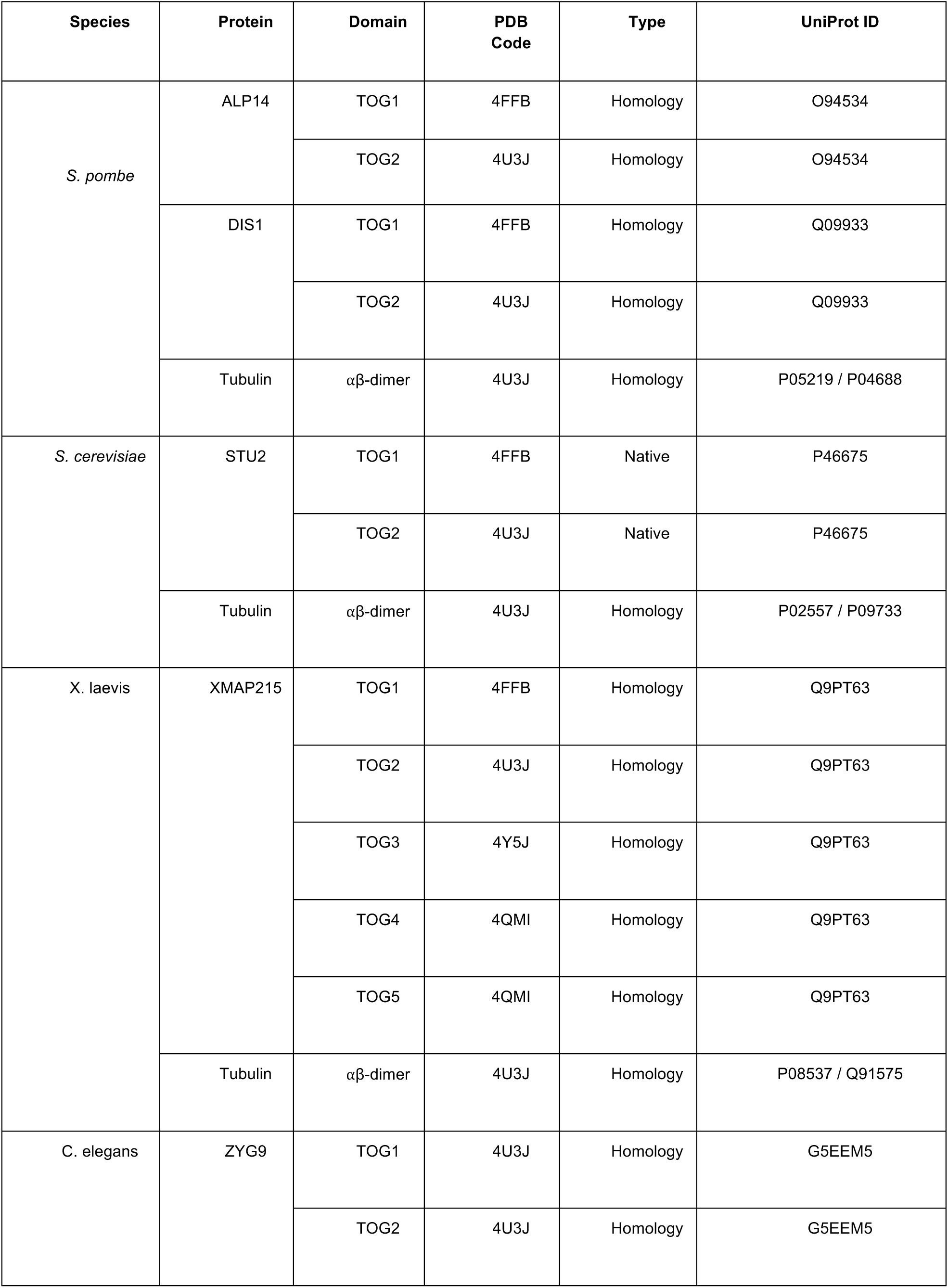
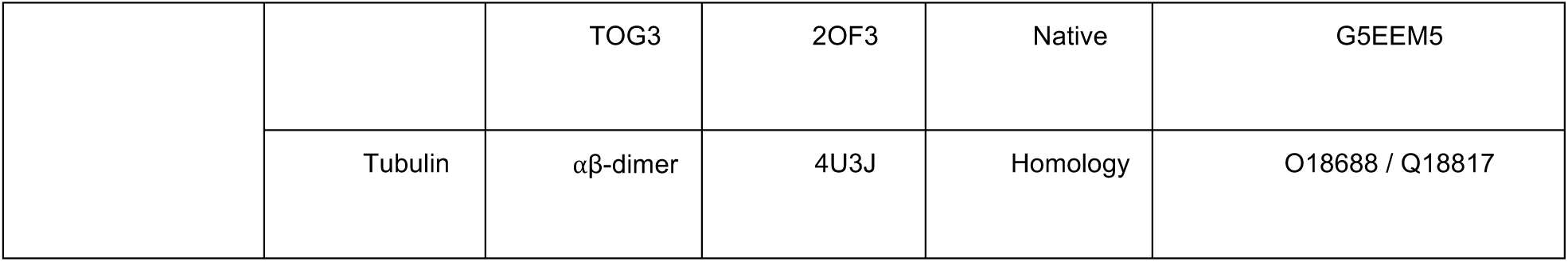
List of structures selected as templates for homology modelling.

## Acknowledgements

Special thanks to Jack Tuszynski for denoting the tubulin C-termini structures.

## Author Contributions

Conceived and designed the experiments: NV RC. Performed and analysed the data: NV.

Wrote the manuscript: NV RC TB

